# CD38–NAMPT–p21 signaling promotes stress adaptation in CLL

**DOI:** 10.64898/2026.11.16.766865

**Authors:** Bing Han, Claudia Chini, Gina L. Ciccio Lopez, Yuki Hamada, Aneel Paulus, Asher Chanan-Khan, Esteban Braggio, Anders Berglund, Eduardo Chini

## Abstract

Chronic lymphocytic leukemia (CLL) is the most common adult leukemia in the United States, accounting for ∼30% of new diagnoses and ∼4,000 deaths annually. High CD38 expression is a well-established marker of poor progno-sis, yet the mechanistic basis of CD38-mediated treatment failure has remained incompletely understood. Here, we demonstrate that CD38 functions not merely as a prognostic biomarker but as a metabolic and signaling regulator that equips CLL cells to withstand cytotoxic and hypoxic stress. Using MEC-1 cells engineered to express CD38, we identified a CD38–NAMPT–p21 survival axis that confers resistance to fludarabine and hypoxia. CD38 NADase activity promoted NAMPT upregulation and AKT activation and was associated with accumulation of cytoplasmic p21. This CD38-dependent phenotype was lost in cells expressing a catalytically inactive CD38 mutant and was also reversed by pharmacologic inhibition of NAMPT (FK866) or suppression of p21 (UC2288). Analysis of three independent single-cell RNA-sequencing datasets of primary CLL further showed that naturally occurring CD38-positive CLL cells exhibit increased NAMPT expression and related stress-response signaling. Analysis of larger primary CLL cohorts further linked NAMPT expression to AKT and hypoxia signatures, providing additional sup-port for the relevance of this pathway in patient-derived CLL. Together, these findings identify CD38–NAMPT– p21 signaling as an important mechanism of stress adaptation in CLL and suggest that targeting NAD⁺ metabolism and p21-mediated survival pathways may represent a potential strategy for CD38-high disease.

**Simple Summary:** Chronic lymphocytic leukemia (CLL) is the most common leukemia in adults, and treatment resistance remains an important clinical challenge. High CD38 expression is associated with more aggressive disease and poorer out-comes, but the mechanisms by which CD38 supports leukemia-cell survival are not fully understood. In this study, we found that CD38 helps CLL cells withstand chemotherapy and low-oxygen stress by altering cellular metabo-lism and promoting survival signals involving NAMPT and p21. Inhibiting NAMPT or p21 reduced this protective effect and restored treatment sensitivity. Analyses of patient CLL datasets showed similar associations between CD38, NAMPT, and stress-related cellular programs. These findings identify a potential mechanism by which CD38 promotes CLL-cell survival and suggest new therapeutic opportunities for CD38-high disease.

## 1. Introduction

Chronic lymphocytic leukemia (CLL) is one of the most common forms of leukemia in the United States, ac-counting for approximately 30% of newly diagnosed leukemia cases and about 1% of all cancers. While most CLL are indolent, the disease remains incurable and causes approximately 4,000 deaths annually [1]. Although advances in therapy have improved the five-year overall survival rate to over 80%, a subset of patients—particularly those with fludarabine-refractory disease—continues to experience poor outcomes and resistance to treatment [2].

Age is a major prognostic factor in CLL, as reflected in the CLL International Prognostic Index (CLL-IPI). According to SEER data, five-year survival declines from nearly 95% in patients younger than 65 years to approximately 85% in those aged 65 and older [3]. This disparity suggests that age-related biological changes—such as cellular senescence, immune dysregulation, and altered metabolism—may drive disease progression and therapeutic resistance.

Among the molecular pathways linking aging and cancer, nicotinamide adenine dinucleotide (NAD⁺) metabolism plays a pivotal role. Maintaining NAD⁺ homeostasis is crucial for cellular metabolism and genome maintenance; however, NAD levels decline during aging and age-related diseases [4–8]. NAD balance depends on the interplay between its synthesis and degradation. Regarding NAD synthesis, the NAD⁺ salvage pathway enzyme nicotinamide phosphoribosyltransferase (NAMPT) is critical and has been found to be dysregulated in many cancers [9–11]. A key mediator of age-related NAD decline is the Cluster of Differentiation 38 (CD38), a multifunctional transmembrane ectoenzyme and receptor that is a major NAD consumer [12, 13]. Through its NADase activity, it regulates NAD⁺ availability while generating second messengers, such as adenosine diphosphate ribose (ADPR) and cyclic ADP-ribose (cADPR), which influence calcium signaling and mitochondrial function [14]. CD38 expression significantly increases with age, especially in pro-inflammatory immune cells, accelerating tissue NAD⁺ consumption, suppressing Poly(ADP-ribose) polymerase (PARP) and sirtuin activity, and promoting cellular and tissue dysfunction [4, 5, 7, 8, 12, 15].

Elevated CD38 expression is frequently observed in CLL and consistently correlates with aggressive disease features, earlier treatment requirement, and poor survival outcomes [16–18]. Beyond serving as a prognostic biomarker, CD38 actively contributes to disease progression by promoting proliferation [19], enhancing survival signaling pathways [20], and facilitating interactions with the tumor microenvironment [21]. Mechanistically, CD38 associates with CD19 and CD81 within lipid rafts, amplifying B cell receptor (BCR) signaling to drive CLL cell activation and growth [22]. Additionally, CD38 interaction with its ligand CD31 promotes signaling programs associated with CLL-cell proliferation, migration, and homing to growth-permissive microenvironments [23]. Clinically, targeting CD38 with monoclonal antibodies has shown promising results in CLL [24, 25], underscoring CD38 as an attractive therapeutic target. However, despite these therapeutic advances, the mechanistic link between CD38’s enzymatic function and treatment resistance remains poorly understood, representing a critical gap in understanding how CD38 shapes therapeutic response and disease progression in CLL.

In the present study, we examined the role of CD38 and its catalytic activity in CLL using engineered MEC-1 cells together with transcriptomic analyses of primary patient samples. We investigated how CD38 NADase activity influences adaptation to fludarabine and hypoxic stress, with particular emphasis on NAMPT, AKT, and p21 signaling. We further evaluated the relevance of this program in primary CLL by examining CD38-associated molecular changes across cohorts, within individual patients, and in the CD38^−^ compartment of patients with differing CD38^+^ cell abundance. Finally, pathway-level and independent large-cohort transcriptomic analyses were used to evaluate the broader relevance of these CD38-associated molecular relationships in primary CLL.

## 2. Materials and Methods

### CD38^WT^ and CD38^CI^ Expression

GeneCopoeia Lenti-Pac HIV Expression Packing Kit (LT002) was used to overexpress wild-type (Gene-Copoeia, EX-G0156-LV207) and catalytically inactive (GeneCopoeia, CS-G0156-LV207-1) CD38 into MEC-1 cells. One day before transfection, 2 x10^6^ of the 293T packaging cells were plated in a 10-cm dish in 10 ml of DMEM with 10% FBS and incubated at 37°C with 5% CO_2_. The day of transfection, 5.0 µl (0.5 µg/µl) of Lenti-Pac HIV Mix and 2.5 µg of lentiviral CD38, CD38^CI^ or control expression plasmid, respectively, were diluted into 200 µl of Opti-MEM® I (Invitrogen) in 5ml sterile polypropylene tubes. In a separate tube, 15 µl of EndoFectin Lenti was diluted into 200 µl of Opti-MEM and then added to EndoFectin Lenti reagent drop-wise to the DNA solution while gently vortexing the DNA-containing tubes. The mixture was incubated for 25 minutes at room temperature to allow the DNA-EndoFectin complex to form. Then the DNA-EndoFectin Lenti complex was added directly to each 293T cell plate, and the plates were gently swirled to distribute the complex. The cells were incubated in a CO_2_ incubator at 37°C overnight. The medium was replaced with fresh DMEM medium with 5% FBS and cultured for additional 48 hours. The pseudovirus-containing culture medium was collected in sterile capped tubes and centrifuged at 500 x g for 10 minutes to eliminate cell debris. Following centrifugation, the supernatant was filtered through 0.45 µm Corning filters. The supernatant containing lentiviral particles was diluted with RPMI1640 medium with 10% FBS at 1:1 ratio and then the Polybrene Infection /Transfection Reagent (SIGMA, TR-1003-G) at 10 μg/mL final concentration was added. 2 mL/well diluted lentivirus was applied on MEC-1 cells and plated in 6-well plate with about 70% confluency. The plate was centrifuged at 1800 rpm room temperature for 1 hour before returning to the 37°C CO_2_ incubator. The virus-containing medium was replaced with regular RPMI 1640 medium 24 hours after and then continued incubation for additional 48 hours before function tests or selection by 200 ug/mL Hygromycin to establish stable cell lines.

### RT-qPCR

Cell samples were harvested and RNA was extracted using Trizol according to the manufacturer’s protocol. Reverse transcription was performed by Bio-Rad iScript supermix with 100 ng RNA per sample. qPCR was run on the Bio-Rad CFX Opus 384 system with Bio-Rad CFX Maestro and Taqman predesigned probe primers (ThermoFisher). Data were analyzed using Bio-Rad CFX Maestro software.

### Cell Culture and Viability Assay

Vector Control, CD38^WT^ and CD38^CI^ MEC-1 cells were cultured by RPMI 1640 medium with 25 mM HEPES and 10% FBS in 37°C 5% CO_2_ 21% O_2_ (normoxia) or 2.5% O_2_ (hypoxia) incubator. Mycoplasma contamination was regularly tested.

For cell growth curve assay, 1,000 cells were seeded per well in 200 µL of culture medium in a 96-well plate. 15 µL of CCK8 cell viability assay (GLPBIO) was added to each well at day 0, 1, 3, 5 and 7. The plate was read at OD450 45 min later.

For drug dose-response assay, 5,000 cells were seeded per well in 200 µL of medium containing the designated drugs, antibodies, or chemicals in a 96-well plate. CCK8 assay was performed 3 days afterwards.

### NADase Activity and Intracellular NAD⁺ Measurement

CD38 NADase activity was measured using the fluorescent substrate 1,N⁶-ethenonicotinamide adenine dinucleotide (ε-NAD), as previously described [26]. Intracellular NAD⁺ levels were measured using an enzymatic cycling assay as previously described [7].

### Subcellular Fraction and Mitochondria Isolation

Pierce Subcellular Protein Fractionation Kit and Mitochondria Isolation Kit were used to isolate subcellular components of cell samples according to manufactory instruction. 20ug of subcellular fraction samples for each lane of immunoblotting.

### Immunoblotting

Cells were washed twice with ice-cold PBS and lysed in RIPA buffer (25 mM Tris-HCl, pH 7.4; 150 mM NaCl; 1 mM EDTA; 1% NP-40; 5% glycerol) supplemented with protease and phosphatase inhibitor cocktails (Roche). Lysates were incubated on ice for 30 minutes and clarified by centrifugation at 14,000 × g for 5 minutes at 4 °C. Protein concentrations were determined using the Bio-Rad Protein Assay, and equal amounts of protein (20–40 µg per sample) were resolved by SDS-PAGE on 10–12.5% polyacrylamide gels and subsequently transferred onto PVDF membranes using a semi-dry transfer system (Bio-Rad). Membranes were blocked for 1 hour at room temperature in 5% non-fat dry milk prepared in TBS-T (20 mM Tris-HCl, pH 7.6; 137 mM NaCl; 0.1% Tween-20) and incubated overnight at 4 °C with primary antibodies diluted in 5% milk/TBS-T. The following antibodies were used: CD38 (Abcam, ab108403), pan-Akt (Cell Signaling Technology, 4691), phospho-Akt (Ser473) (Cell Signaling Technology, 4060), GAPDH (Cell Signaling Technology, 97166S), β-Tubulin (Abcam, ab15558), Cox IV (Cell Signaling Technology, 4850S), p21Waf1/Cip1 (Cell Signaling Technology, 37543), Caspase-3 (Cell Signaling Technology, 9662S), and Bcl-xL (Cell Signaling Technology, 2764S). After washing, membranes were incubated with HRP-conjugated secondary antibodies (Cell Signaling Technology) diluted 1:5000 in 5% non-fat milk/TBS-T for 1 hour at room temperature. Protein bands were detected using the SuperSignal™ West Pico PLUS chemiluminescent substrate (Thermo Scientific, 34580) and visualized with the Amersham Imager 600 system. Images were processed using ImageJ. Graphs show protein quantification normalized to tubulin or GAPDH levels.

### Data Analysis of CLL Patient Samples from GEO datasets

#### Single-cell RNA-sequencing analysis of primary CLL

Publicly available scRNA-seq datasets GSE111014, GSE165087, and GSE259253 were obtained from the Gene Expression Omnibus [27–29]. For GSE111014, only baseline (day 0) samples were included. Single-cell data processing and visualization were performed using Scanpy [30]. Cells were retained using quality-control thresholds of 300–2500 detected genes, at least 500 total counts, and <15% mitochondrial reads. The B/CLL compartment was identified using relative lineage scores derived from the B-cell markers CD79A, CD79B, MS4A1, CD37, CD22, HLA-DRA, CD19, and CD74, in comparison with T-cell, NK-cell, and myeloid marker sets. Within the B/CLL compartment, CD38^+^ and CD38^−^ groups were defined by detectable and undetectable CD38 transcript counts, respectively, before normalization; these designations therefore refer to transcript detection rather than flow-cytometric surface CD38 expression. Counts were library-size normalized to 10,000 counts per cell and log transformed. One GSE165087 sample, CLL7, was excluded from all final analyses and visualizations because cohort-level quality-control review identified an anomalous reconstruction/QC profile; all GSE165087 results were subsequently regenerated after exclusion of this sample. The retained datasets comprised 4, 8, and 6 patients from GSE111014, GSE165087, and GSE259253, respectively. The two GSE165087 CLL1 technical libraries were combined for patient-level analyses.

#### CD38^+^ versus CD38^−^ gene-level analyses

Gene-level comparisons focused on NAMPT, HIF1A, and CDKN1A. For the cohort-balanced summary analysis, the percentage of cells with detectable transcript and the mean normalized expression within each CD38 group were calculated separately for each cohort and then averaged with equal weight across GSE111014, GSE165087, and GSE259253. Expression-distribution comparisons were evaluated using Mann–Whitney tests, and detection-frequency differences were evaluated using Fisher’s exact tests. For cross-cohort inference, cohort-specific 2 × 2 detection tables were combined using dataset-stratified Mantel–Haenszel (MH) odds ratios (ORs). False-discovery-rate (FDR) correction was applied to the gene-detection analyses, and Breslow– Day tests were used to assess heterogeneity among cohorts.

To evaluate the CD38–NAMPT association within individual patients, NAMPT-detection tables were constructed separately for each patient, allowing direct comparison of CD38^+^ and CD38^−^ cells from the same disease sample. Only patients containing cells in both CD38 groups contributed an estimable within-patient OR; GSE165087 CLL8 and GSE259253 Pt5 contained no CD38^+^ cells and therefore did not contribute to this analysis. This yielded 4, 7, and 5 evaluable patients from GSE111014, GSE165087, and GSE259253, respectively, for a total of 16 patients. For patient-specific ORs and normal-approximation confidence intervals, 0.5 was added to all four cells of a 2 × 2 table when a zero count was present; Fisher’s exact tests were performed using the original counts. Cohort-specific and overall patient-stratified MH estimates were calculated from the uncorrected patient tables. The overall estimate was obtained directly across all 16 patient strata, rather than by pooling the three cohort-level ORs using random-effects meta-analysis.

#### Patient-context analysis of the CD38^−^ compartment

Relative CD38^+^ abundance was defined for each patient as the percentage of B/CLL cells with detectable CD38 transcript. NAMPT detection within the CD38^−^ compartment was defined as the percentage of CD38^−^ B/CLL cells with detectable NAMPT transcript. To account for differences among datasets, each variable was ranked within its respective cohort, divided by the number of patients in that cohort, and expressed on a 0–100 scale. Spearman rank correlation between relative CD38^+^ abundance and NAMPT detection in the CD38^−^ compartment was then calculated across the 18 retained patients. FDR correction was applied across seven prespecified CD38^−^-compartment outcomes: detection frequencies of NAMPT, HIF1A, and CDKN1A and mean scores for Hypoxia, PI3K/AKT/mTOR, mTORC1, and Glycolysis. Patients without detectable CD38 transcript were retained because their CD38^−^ compartment remained evaluable. The fitted line shown in Figure 7f is provided as a visual guide; statistical inference was based on the Spearman rank correlation.

#### Pathway-score analysis

Hypoxia, PI3K/AKT/mTOR, mTORC1, and Glycolysis gene sets were obtained from the MSigDB Hallmark collection [31]. Pathway scores were calculated within each patient’s B/CLL population using the Scanpy score_genes function, genes present in the expression matrix, and a fixed random seed of 42. Patients were eligible for pathway effect-size analysis when at least five CD38^+^ B/CLL cells were available. After application of the quality-control exclusion described above, 4 patients from GSE111014, 6 from GSE165087, and 4 from GSE259253 contributed to the analysis (14 patients in total); GSE259253 Pt2 met the eligibility threshold and was included. Within each patient, Hedges’ g was calculated as the difference between the mean pathway scores of CD38^+^ and CD38^−^ cells divided by the pooled within-group standard deviation and multiplied by the Hedges small-sample correction. Positive Hedges’ g therefore indicates a higher pathway score in CD38^+^ cells. Patient-level effect estimates were pooled within each cohort using inverse-variance random-effects meta-analysis with DerSimonian– Laird estimation of τ², and cohort-level estimates were subsequently combined using the same random-effects procedure. Normal-approximation 95% confidence intervals and two-sided P values were calculated, and Cochran’s Q and I² were used to characterize heterogeneity. Hedges’ g represents a standardized difference in pathway score rather than a direct measurement of pathway activation.

#### Single-cell visualization

UMAP embeddings were generated from highly variable genes following principal-component analysis and nearest-neighbor graph construction. CD38 and NAMPT were excluded from the highly variable gene set used to generate embeddings.

#### Large-cohort transcriptomic analysis

Raw CEL files from GSE58211 (n = 300) were downloaded from GEO and normalized using IRON [32] with the Version 25 ENTREZ gene CDF from BrainArray [33]. Updated gene annotation (Homo_sapiens.gene_info) was obtained from NCBI. For GSE246521 (n = 350), raw RNA-seq gene counts were obtained from the corresponding author [34], normalized using DESeq2, and batch adjusted using COMBAT. Hypoxia and AKT gene sets were obtained from the MSigDB Hallmark collection [31]. HYPOXIA and AKT scores were calculated from the first principal component of a principal-component analysis using the genes comprising each respective signature [35]. Associations of NAMPT expression with AKT Score, HYPOXIA Score, and CDKN1A expression were evaluated separately within each dataset using Pearson correlation.

## 3. Results

### 3.1. Expression of CD38 in MEC-1 Cells Does Not Alter Cell Proliferation Under Normoxic Conditions

To investigate the contribution of CD38 to the biology of CLL, we generated a CD38-expressing MEC-1 cell line (CD38^WT^) via lentiviral transduction of the human *CD38* gene under control of a CMV promoter. Following infection and antibiotic selection, three independent clones were established and selected for further characterization (Figure 1a). To confirm that CD38 expression resulted in functionally active protein, we measured CD38 enzymatic activity. All CD38^WT^ clones displayed robust NADase activity, consistent with the canonical role of CD38 as a major NAD⁺ hydrolase (Figure 1b, pooled clones; and Supplementary Figure S1a, individual clones).

Given the cell type–specific subcellular localization of CD38 [36–40], we next examined CD38 subcellular distribution in MEC-1 cells. Flow cytometry demonstrated robust surface expression of CD38 (Figure 1c, pooled clones; Supplementary Figure S1b, representative flow cytometry plot), whereas subcellular fractionation followed by immunoblotting revealed its presence in the mitochondrial compartment, but not in the cytosol (Figure 1d). This dual localization suggests that CD38 may participate in both extracellular and organelle-specific NAD⁺ metabolism.

To assess whether CD38 expression impacts cell proliferation, we compared the growth kinetics of CD38^WT^ and vector-transduced control cells. At low seeding density, CD38^WT^ cells exhibited a modest delay in proliferation. Interestingly, as cell density increased, the proliferation of CD38^WT^ cells accelerated and eventually became indistinguishable from control cells (Figure 1e, representative clones; Supplementary Figure S1c, all tested clones). These findings suggest that CD38 does not impair long-term cell growth under normoxic conditions, but it may impose a transient metabolic burden at early stages of cell culture.

**Figure 1.**
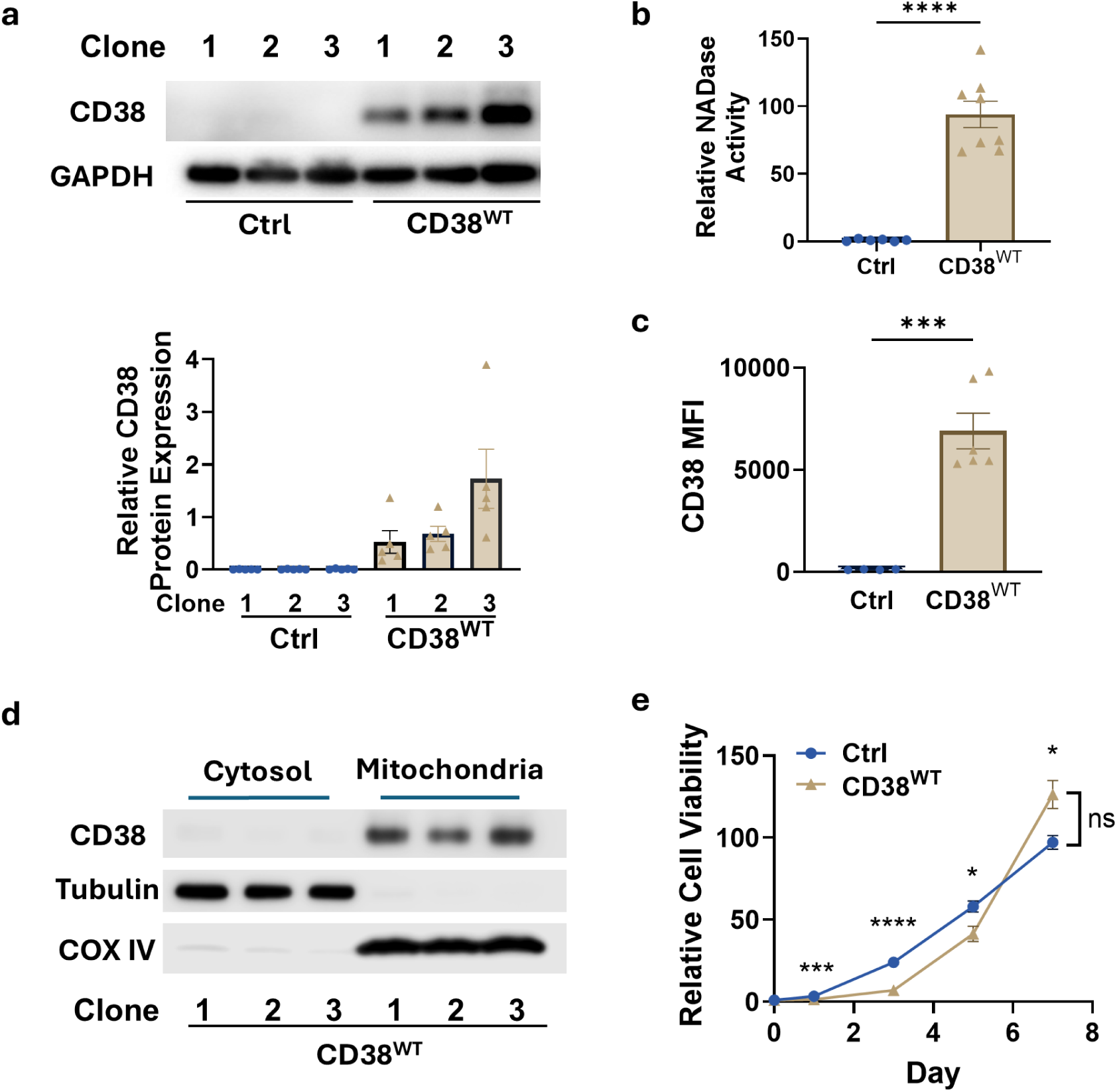
CD38 Overexpression in MEC-1 Cells. (a) Immunoblot analysis of CD38 expression in three independent clones of vector control and CD38-overexpressing (CD38^WT^) MEC-1 cells. Semi-quantification from five independent experiments is shown below; (b) ε-NAD fluorescence–based NADase assay demonstrating catalytic activity of CD38^WT^ cells. Quantification of relative enzymatic activity from pooled clones is shown (control, n=6; CD38^WT^, n=8; unpaired *t*-test ****: p<0.0001); (c) Flow cytometry analysis of surface CD38 expression in vector control and CD38^WT^ cells. Mean fluorescence intensity (MFI) values from multiple clones are summarized (control, n=4; CD38^WT^, n=6; unpaired *t*-test ***: p<0.001); (d) Immunoblotting of subcellular fractions showing localization of CD38 in the mitochondrial but not cytosolic fraction of CD38^WT^ cells. Representative results from three independent clones are shown; (e) Growth curves of vector control and CD38^WT^ cells demonstrating comparable cell proliferation. Results from three independent experiments of representative clones are shown; data are presented as fold change relative to Day 0 and analyzed by two-way ANOVA and unpaired *t*-test ( *: p < 0.05, **: p < 0.01, ***: p < 0.001, ****: p<0.0001).

### 3.2. CD38 Expression in MEC-1 Cells Confers Hypoxia Tolerance and Fludarabine Resistance

Although CD38 had little effect on cell growth under normoxic conditions, its impact became pronounced under stress. When cultured in hypoxia (2.5% O₂), vector control MEC-1 cells showed profound growth inhibition, whereas CD38^WT^ cells maintained stable proliferation rates (Figure 2a, representative clones; Supplementary Figure S1d, all tested clones). To uncover the molecular basis of this adaptation to limited oxygen, we analyzed the expression of key hypoxia-responsive genes. Compared to control cells, CD38^WT^ cells exhibited robust induction of *HIF1A*, *VEGFA*, *LDHA*, and *SLC2A1*/*GLUT1* mRNAs under hypoxic conditions (Figure 2b, representative clones; Supplementary Figure S1e, all tested clones). These genes represent crucial components of the HIF-1 transcriptional program: angiogenesis (*VEGFA*), glycolytic metabolism (*LDHA*), and glucose uptake (*SLC2A1*/*GLUT1*). Their upregulation suggests that CD38 enhances hypoxia signaling and promotes metabolic reprogramming toward glycolysis, supporting cellular adaptation to oxygen deprivation. Importantly, these findings connect CD38 activity to key metabolic pathways involved in cellular stress responses.

We next assessed whether CD38 also impacts chemotherapeutic sensitivity, given the clinical association between CD38 expression and poor prognosis in CLL. Vector control cells and CD38^WT^ cells were treated with increasing concentrations of fludarabine, a frontline nucleoside analog in CLL therapy that induces apoptosis. CD38^WT^ cells displayed an extraordinary increase in resistance to fludarabine, with an approximately 1800-fold higher IC₅₀ relative to control cells (Figure 2c). This profound drug resistance was observed without previous drug exposure, indicating that CD38 expression confers intrinsic resistance, rather than being acquired through prolonged drug exposure.

To further characterize this phenotype, we measured apoptosis following fludarabine treatment or exposure to severe hypoxia (1% O₂) using Annexin V/PI staining. In both conditions, CD38^WT^ cells exhibited significantly lower levels of early apoptotic and late-apoptotic/dead cells than control cells (Figure 2d, representative clones; Supplementary Figure S1f, cell death of all tested clones). These results confirm that CD38 protects MEC-1 cells from both genotoxic and metabolic stress, extending beyond its effects on proliferation under hypoxia.

**Figure 2.**
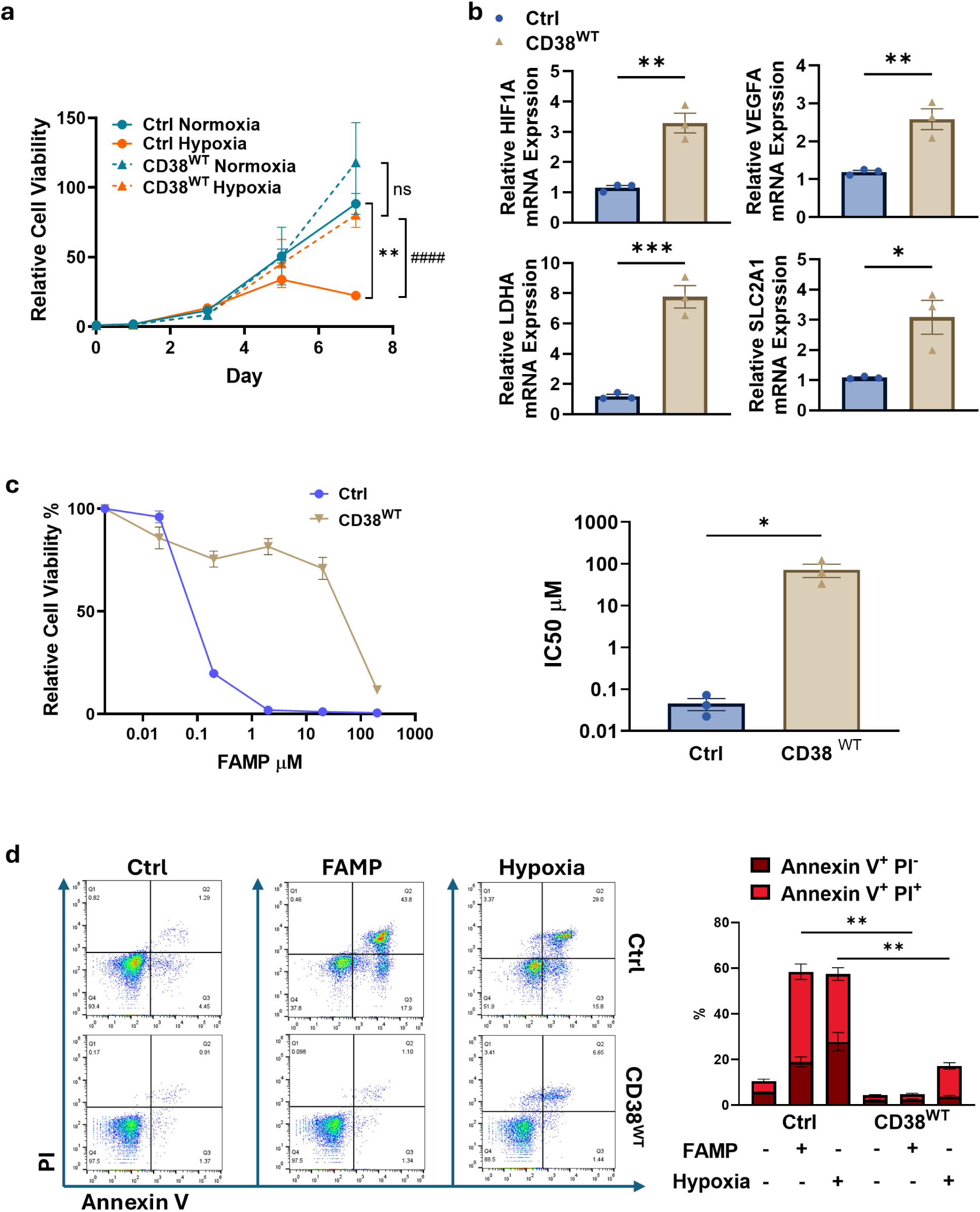
CD38 Expressed MEC-1 Cells Gain Hypoxia Tolerance and Fludarabine Resistance. (a) Proliferation curves of representative vector control and CD38^WT^ clones cultured under normoxic and hypoxic conditions, data are presented as fold change relative to Day 0 (n=3; two-way ANOVA, **: p < 0.01, unpaired *t*-test, ####: p < 0.0001); (b) Quantitative RT–PCR analysis of hypoxia-responsive gene expression in vector control and CD38^WT^ cells following hypoxic exposure (2.5% O2). Results from a representative clone are shown (n=3; unpaired *t*-test, *: p < 0.05, **: p < 0.01, ***: p < 0.001); (c) Fludarabine (FAMP) dose–response curves after 72 hours of treatment (left) and corresponding mean IC₅₀ values (right) comparing vector control and CD38^WT^ cells. Average of different clones from three independent experiments are shown (unpaired *t*-test, *: p < 0.05); (d) Flow cytometry analysis of apop tosis following 72 hours of fludarabine treatment or hypoxia exposure (1% O2). Representative Annexin V/PI plots are shown (left), with quantification of early apoptotic cells (Annexin V⁺ PI⁻) and late-apoptotic/dead cells (Annexin V⁺ PI⁺) across representative clones (right, n=4; unpaired *t*-test, **: p < 0.01).

### 3.3. CD38 Induces Cytoplasmic p21 and Anti-Apoptotic Signaling

CD38 has been linked to altered cellular metabolism, proliferation, survival, and tumor progression in several cancer contexts, including CLL [19], cervical cancer [41], breast cancer [42], and ovarian cancer [43]. In our model, CD38-expressing MEC-1 cells (CD38^WT^) showed marked upregulation of p21Cip1/Waf1 (CDKN1A) compared to vector control cells (Figure 3a, protein levels by immunoblotting and pooled mRNA levels from all tested clones). This finding is consistent with previous studies in pancreatic cancer [9] and prostate cancer [44], in which experimental CD38 expression was associated with increased p21 expression. However, despite elevated p21 expression—a classic marker of cell cycle arrest— CD38^WT^ cells exhibited no proliferative defects and instead continued to divide robustly (Figure 2a). These findings suggest a non-canonical role for p21, independent of its traditional function as a cyclin-dependent kinase inhibitor. To explore this possibility, we performed subcellular fractionation and immunoblot for p21 in CD38^WT^ cells. This experiment revealed that p21 is predominantly localized to the cytoplasm in these cells (Figure 3b), in contrast to the nuclear localization typically associated with cell growth arrest [45].

Cytoplasmic p21 can promote apoptosis resistance by suppressing caspase-3 activation, including through direct interaction with procaspase-3 [46, 47]. Indeed, CD38^WT^ cells treated with fludarabine exhibited markedly reduced caspase-3 cleavage (CC3) compared to control cells (Figure 3c, left and middle panels), indicating suppressed apoptotic signaling. Under hypoxic stress, which mimics the bone marrow and lymph node microenviron-ments and imposes metabolic and oxidative pressure, caspase-3 cleavage was low in both CD38^WT^ and control cells (Figure 3c, left and middle panels). This observation suggests that fludarabine and hypoxia may engage distinct cell death pathways, with the latter likely invoking mostly caspase-independent mechanisms. Notably, CD38^WT^ cells maintained higher levels of B-cell lymphoma-extra large (Bcl-xL), a key anti-apoptotic member of the B-cell lymphoma 2 (BCL-2) family, than control cells under both stress conditions (Figure 3c, left and right panels). Given the established role of p21 in promoting resistance to stress-induced apoptosis, including through Bcl-xL–dependent mechanisms [48, 49], these findings support a model in which CD38-associated p21 upregulation contributes to cell survival through maintenance of Bcl-xL expression and suppression of caspase activity.

To assess the functional importance of p21 in this adaptive survival program, we treated CD38^WT^ cells with UC2288, a relatively selective small-molecule p21 attenuator [50]. UC2288 markedly reduced *p21*/*CDKN1A* mRNA and protein expression (Figure 3d, pooled protein levels from all tested clones by immunoblotting and pooled mRNA levels from all tested clones), significantly increased the sensitivity of CD38^WT^ cells to fludarabine-induced cell death (Figure 3e) and suppressed cell growth in hypoxic conditions (Figure 3f). This was accompanied by decreased expression of hypoxia-responsive genes, including *HIF1A*, *VEGFA*, *LDHA* and *SLC2A1*/*GLUT1* (Figure 3g). These results suggest that p21 not only contributes to apoptosis resistance but also supports metabolic and hypoxic adaptations downstream of CD38. Together, these findings establish a mechanistic link between CD38, cytoplasmic p21 accumulation, and enhanced anti-apoptotic signaling in CLL cells.

**Figure 3.**
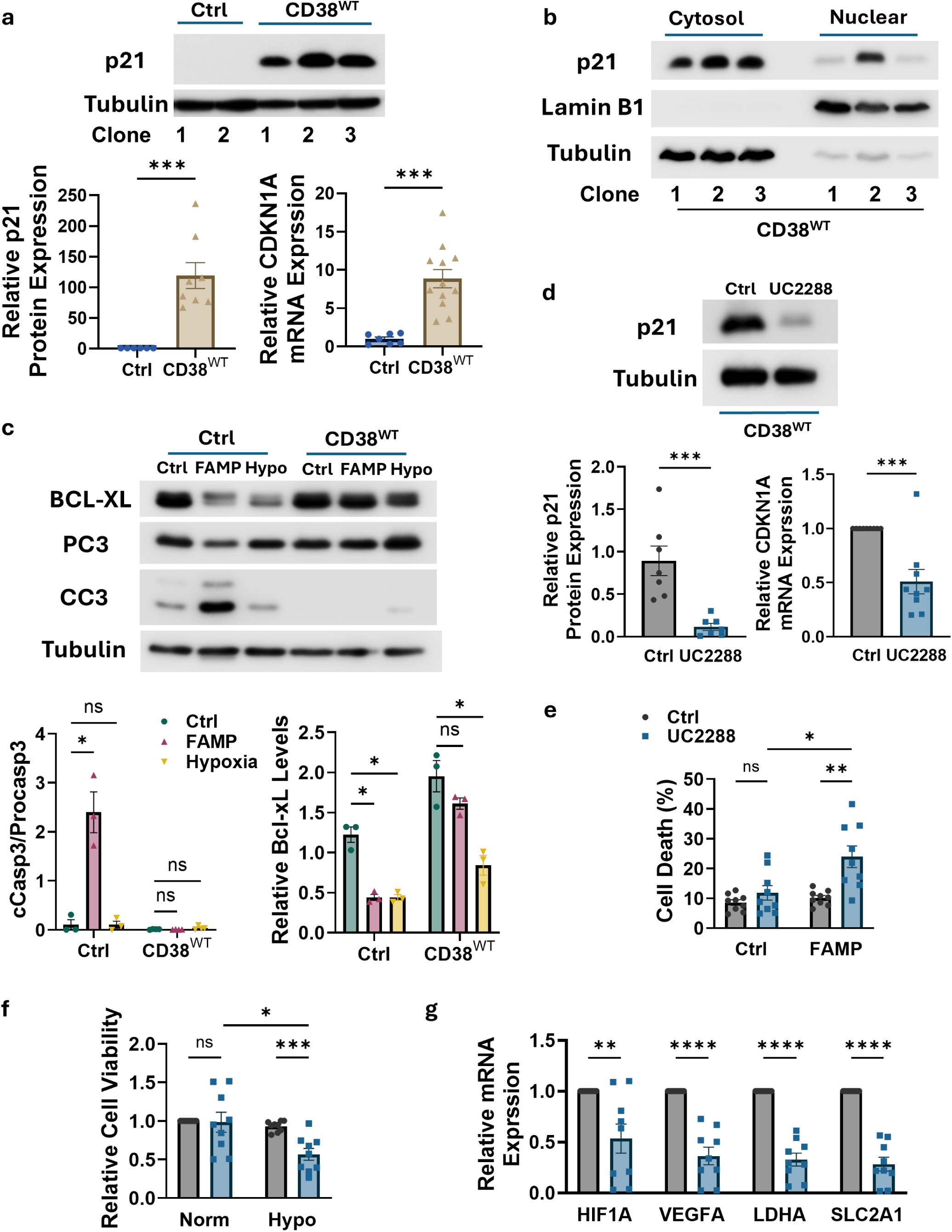
CD38 Induces Cytoplasmic p21 and Anti-Apoptotic Signaling. (a) Immunoblot analysis showing robust upregulation of p21Cip1/Waf1 in CD38^WT^ MEC-1 cells compared with vector control cells. Semi-quantification from multiple clones is summarized below (control, n=6; CD38^WT^, n=9; unpaired *t*-test, ***: p < 0.001); (b) Subcellular fractionation and immunoblot analysis demonstrating predominant cytoplasmic localization of p21 in CD38^WT^ cells. Representative blot from three independent clones is shown; (c) Immunoblot analysis of apoptosis- and survival-associated proteins in response to fludarabine (FAMP) or hypoxic stress (Hypo). CD38^WT^ cells displayed reduced cleaved caspase-3 (CC3, middle panel) and sustained Bcl-xL expression (right panel) compared with controls after 72h treatments. Semi-quantification from representative clones across three independent experiments is shown (unpaired *t*-test, *: p < 0.05); (d) Immunoblot analysis showing that 7 days of treatment with the p21 attenuator UC2288 (5 μM) suppressed p21 expression in CD38^WT^ cells. Representative blot (left) and semi-quantification from multiple clones are shown (n=6; unpaired *t*-test, ***: p < 0.001); (e–f) Pretreatment with UC2288 (5 μM) for 7 days significantly sensitized CD38^WT^ cells to fludarabine-induced cell death (e) and hypoxia-induced growth inhibition (f) after 72h treatments. Pooled results from different CD38^WT^ clones are shown (n=9; unpaired *t*-test, *: p < 0.05, **: p < 0.01); (g) UC2288 treatment diminished induction of hypoxia-responsive genes under hypoxic conditions. Pooled results from different CD38^WT^ clones are shown (n=9; unpaired *t*-test, **: p < 0.01, ****: p < 0.0001).

### 3.4. CD38-Driven Resistance Is Dependent on its NADase Activity

To determine whether the enzymatic activity of CD38 is required for its pro-survival effects, we generated MEC-1 cells expressing catalytically inactive CD38 (CD38^CI^), carrying the E226D substitution [5, 51]. This mutation disrupts CD38’s NADase activity while preserving its membrane localization and potential receptor-like signaling functions, allowing us to separate enzymatic activity from non-catalytic CD38 signaling. For comparisons with CD38^WT^, we selected clones with similar expression levels or combined data from multiple clones. CD38^CI^ expression was confirmed by both immunoblotting (Figure 4a) and flow cytometry (Supplementary Figure S2a), and the loss of enzymatic activity was verified using a NADase assay (Figure 4b).

Compared to CD38^WT^-expressing cells, CD38^CI^-expressing cells showed no protection against stress. Specifically, CD38^CI^ cells were as sensitive to fludarabine-induced cytotoxicity as control cells, showing marked reductions in cell viability (Figure 4c and Supplementary Figure S2b) and they remained highly sensitive to hypoxic stress (Figure 4d, representative clones; Supplementary Figure S2c, all tested clones of CD38^CI^). In addition, they exhibited similar levels of cell death as control cells when exposed to both fludarabine and hypoxia conditions (Figure 4e, representative clones; Supplementary Figure S2d, all tested clones of CD38^CI^). Mechanistically, the loss of NADase activity also abrogated CD38’s ability to induce p21 accumulation (Figure 4f), which is associated with stress adaptation and drug resistance.

Together, these results establish that CD38’s enzymatic activity is necessary for its pro-survival role in MEC-1 CLL cells. The inability of CD38^CI^ to confer resistance indicates that the receptor-like signaling alone is not sufficient. Instead, the metabolic effects of CD38 enzymatic activity—rather than its membrane-associated signaling— serve as the primary driver of CD38-mediated chemoresistance and adaptation to hypoxia.

**Figure 4.**
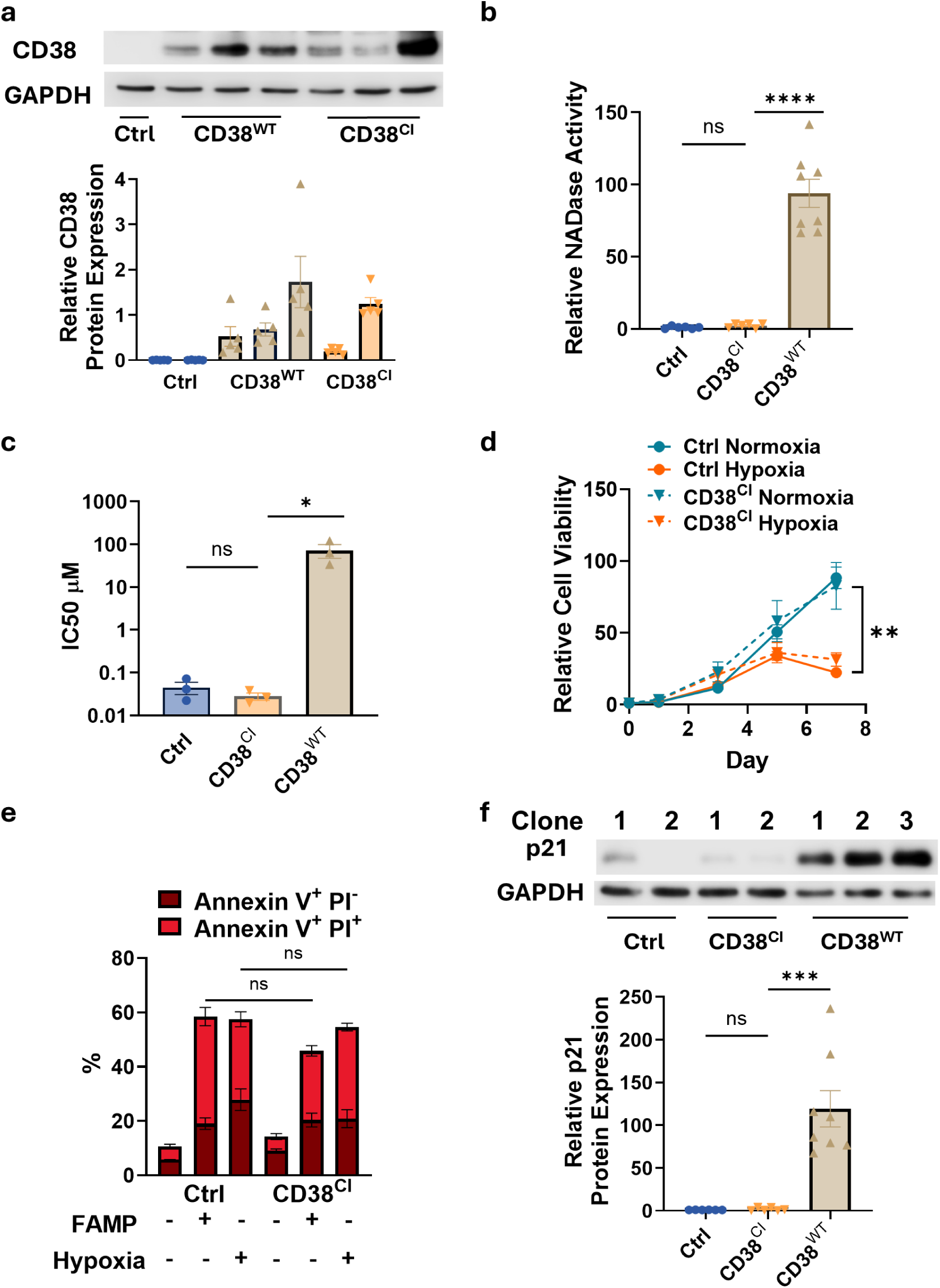
CD38-Driven Resistance depends on its Enzymatic Activity. (a) Generation of CD38^CI^ cells and confirmation of comparable CD38 protein expression in CD38^CI^ and CD38^WT^ clones by immunoblotting. Semi-quantification from five independent experiments is shown below; (b) NADase activity assay demonstrating loss of catalytic ac tivity in CD38^CI^ cells compared with CD38^WT^ cells. Quantification of relative enzymatic activity from pooled clones is shown (control, n=6; CD38^CI^, n=6; CD38^WT^, n=8; unpaired *t*-test, ****: p < 0.0001); (c) Mean IC₅₀ values for fludarabine in vector control, CD38^CI^, and CD38^WT^ cells. CD38^CI^ cells retained sensitivity comparable to vector controls. Data represents averages from multiple clones across three independent experiments (unpaired *t*-test, *: p < 0.05); (d) CD38^CI^ cells remained sensitive to hypoxic stress and failed to acquire the resistance observed in CD38^WT^ cells. Representative growth curves of vector control and CD38^CI^ clones cultured under normoxic and hypoxic con ditions are shown, data are presented as fold change relative to Day 0 (n=3; two-way ANOVA, **: p < 0.01); (e) Quantification of apoptosis following 72 hours of fludarabine or hypoxia treatment showing CD38^CI^ cells behaved similarly to vector controls, in contrast to CD38^WT^ cells (unpaired *t*-test); (f) Immunoblot analysis of p21 expression showing that CD38^CI^ cells did not upregulate p21, indicating a requirement for CD38 catalytic activity in activation of this adaptive survival pathway. Semi-quantification from pooled clones is shown (control, n=6; CD38^CI^, n=6; CD38^WT^, n=9; unpaired *t*-test, ***: p < 0.001).

### 3.5. CD38-Induced Resistance In MEC-1 Cells is Dependent on NAMPT

Despite the high levels of CD38 (Figure 1b), CD38^WT^ cells maintain intracellular NAD^+^ levels comparable to those of CD38^CI^ and control cells (Figure 5a). This unexpected result suggests that CD38 expression may trigger a compensatory upregulation of NAD^+^ biosynthesis in MEC-1 cells. Indeed, among the NAD metabolism–related genes examined (Supplementary Figure S3a), CD38^WT^ cells displayed elevated expression of nicotinamide phosphoribosyltransferase (NAMPT)—the rate-limiting enzyme in the NAD⁺ salvage pathway—compared with CD38^CI^ and vector control cells (Figure 5b representative clones; Supplementary Figure S3b, all tested clones). This suggests that CD38-driven NAD^+^ consumption may trigger NAMPT upregulation, which recycles nicotinamide via the salvage pathway to maintain cellular NAD levels.

To assess the functional relevance of this compensatory response, we treated cells with 5 nM FK866, a highly specific NAMPT inhibitor [52]. CD38^WT^ cells were particularly sensitive to FK866, exhibiting marked reductions in both cell viability and NAD levels, while CD38^CI^ and control cells were less affected (Figure 5c, NAD level of representative clones; Supplementary Figure S3c, cell viability of representative clones). These results support a model in which CD38-mediated NAD^+^ degradation creates a metabolic dependency on NAMPT, establishing a salvage loop that is critical for cell survival under conditions of NAD^+^ consumption.

NAMPT has also been shown to support pro-survival signaling through AKT, with NAMPT inhibition reducing AKT phosphorylation in leukemia cells [53] and NAMPT promoting AKT activation in breast cancer cells [54]. Because AKT can stabilize cytoplasmic p21 [55–58], we next examined AKT activation in CD38^WT^ cells. Compared to CD38^CI^ and control cells, CD38^WT^ cells showed significantly higher levels of phosphorylated AKT (Ser473, p-AKT) (Figure 5d). Importantly, FK866 treatment in CD38^WT^ cells markedly suppressed p-AKT and decreased p21 protein levels (Figure 5e). Moreover, FK866 increased cell death and decreased viability in CD38^WT^ under basal conditions and following exposure to fludarabine or hypoxia (Figure 5f). Collectively, these findings indicate that NAMPT activity is necessary for maintaining AKT signaling and downstream p21-mediated resistance.

**Figure 5.**
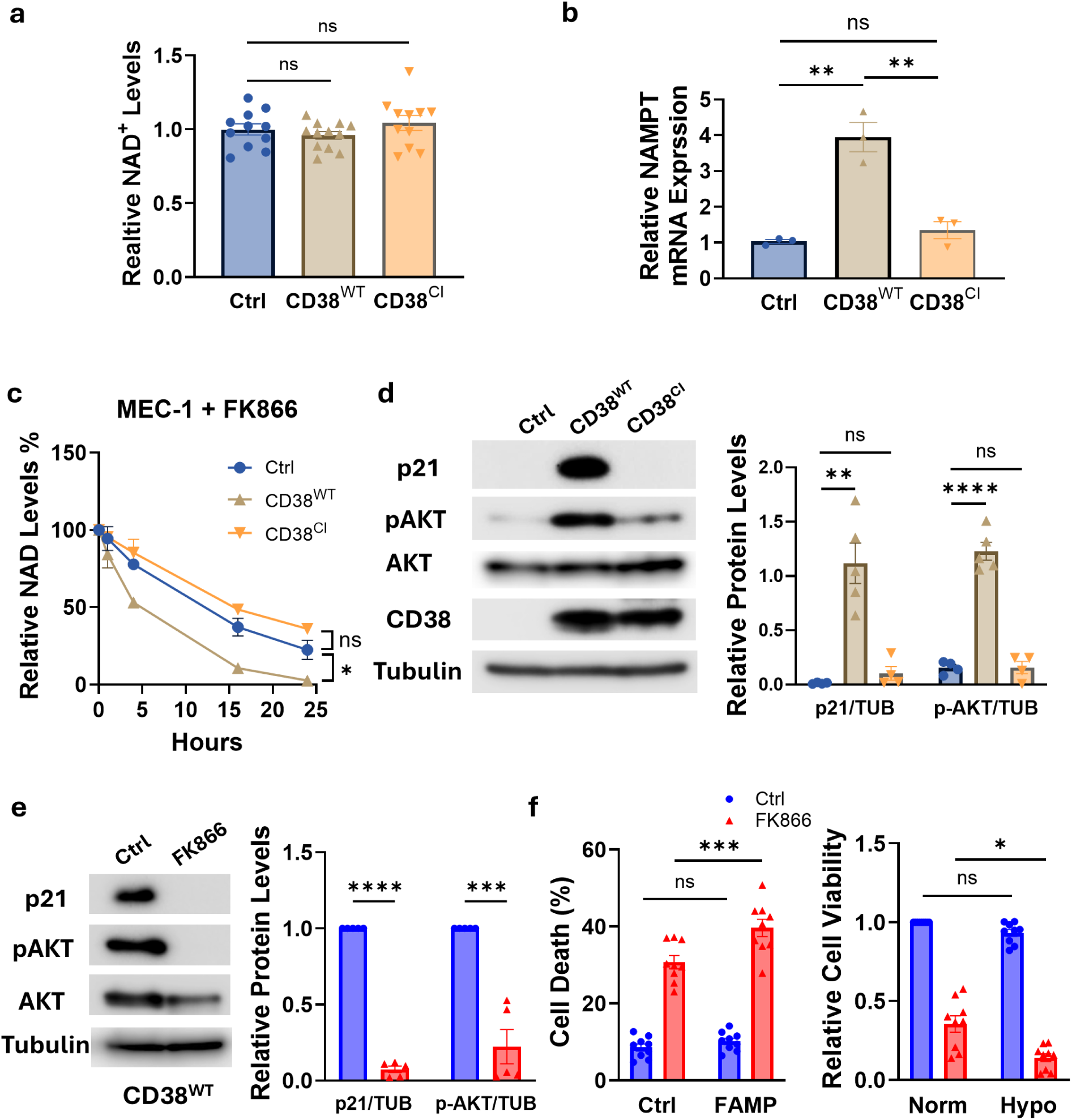
CD38 Driven Resistance depends on NAMPT Activity. (a) Intracellular NAD⁺ levels in CD38^WT^, CD38^CI^, and vector control MEC-1 cells. Quantification from pooled clones is shown (n=11; unpaired *t*-test); (b) Quantitative RT-PCR analysis showing selective upregulation of NAMPT in CD38^WT^ cells, consistent with maintenance of total NAD⁺ despite increased NADase activity. Data represents representative clones across three independent experiments (unpaired *t*-test, **: p < 0.01); (c) CD38^WT^ cells displayed heightened sensitivity to the NAMPT inhibitor FK866 (5 nM), with accelerated NAD⁺ depletion compared with vector control and CD38^CI^ cells. Data represents representative clones across three independent experiments (two-way ANOVA, *: p < 0.05); (d) Immunoblot analysis showing increased AKT phosphorylation and elevated p21 in CD38^WT^ cells relative to vector control and CD38^CI^ cells. Representative blot (left) and semi-quantification from four independent experiments (right) are shown (unpaired *t*-test, **: p < 0.01, ****: p < 0.0001); (e) 96 hours of FK866 (20 nM) treatment suppressed AKT phosphorylation and reduced p21 expression in CD38^WT^ cells. Representative blot (left) and semi-quantification from four independent experiments (right) are shown (unpaired *t*-test, ***: p < 0.001, ****: p < 0.0001); (f) Pretreatment with FK866 (5 nM) for 96 hours restored CD38^WT^ cell sensitivity to fludarabine and hypoxic stress. Fludarabine-induced cell death from pooled clones is shown (left; n=9, unpaired *t*-test, ***: p < 0.001), and hypoxia-induced reduction in cell viability is shown (right; n=9, unpaired *t*-test, *: p < 0.05).

### 3.6. Therapeutic Strategies for CD38^WT^ MEC-1 Cells: Effect of CD38 Inhibitors and Senolytic Agents

In this study, we observed that CD38 promotes resistance to both fludarabine and hypoxia in MEC-1 cells. Besides testing UC2288 and FK866—which target p21 and NAMPT, respectively, to counteract resistance—we also explored the therapeutic potential of CD38-targeting antibodies, enzymatic inhibitors, and senolytic agents.

Daratumumab and Isatuximab are FDA-approved human monoclonal antibodies used to treat multiple myeloma characterized by high CD38 expression [59, 60]. Interestingly, while these antibodies did not affect the viability of CD38^WT^ MEC-1 even at high doses (Figure 6a), they significantly suppressed cell growth under hypoxic conditions, with no significant effect on vector control cells (Figure 6b). This finding suggests a hypoxia-specific vulnerability of CD38^WT^ cells to antibody-based therapies.

We next tested 78c, a potent and specific small-molecule inhibitor of CD38 enzymatic activity [5, 15, 61]. Unlike CD38-targeting antibodies, which act only on cell surface CD38, and are not prominent enzymatic inhibitors, 78c inhibits the enzymatic activity of both surface and intracellular CD38. CD38^WT^ cells showed marked sensitivity to 78c even under normoxic conditions, while control cells were not significantly affected (Figure 6c left panel: normoxia condition; right panel: hypoxia condition 2.5% O2).

Given the prominent role of cytoplasmic p21 in CD38^WT^ cell resistance and its association with senescence, we examined senolytic agents known to target p21^high^ cells. CD38^WT^ cells were markedly sensitive to the BCL-2 family inhibitor Navitoclax (ABT-263), which significantly reduced their viability compared to control cells (Figure 6d). Dasatinib enhanced fludarabine-associated loss of viability in CD38^WT^ cells (Figure 6e, left panel). When tested alone at 0.15 µM, dasatinib reduced relative viability more strongly in CD38^WT^ cells than in vector controls (Figure 6e, right panel), supporting further evaluation of this combination.

**Figure 6.**
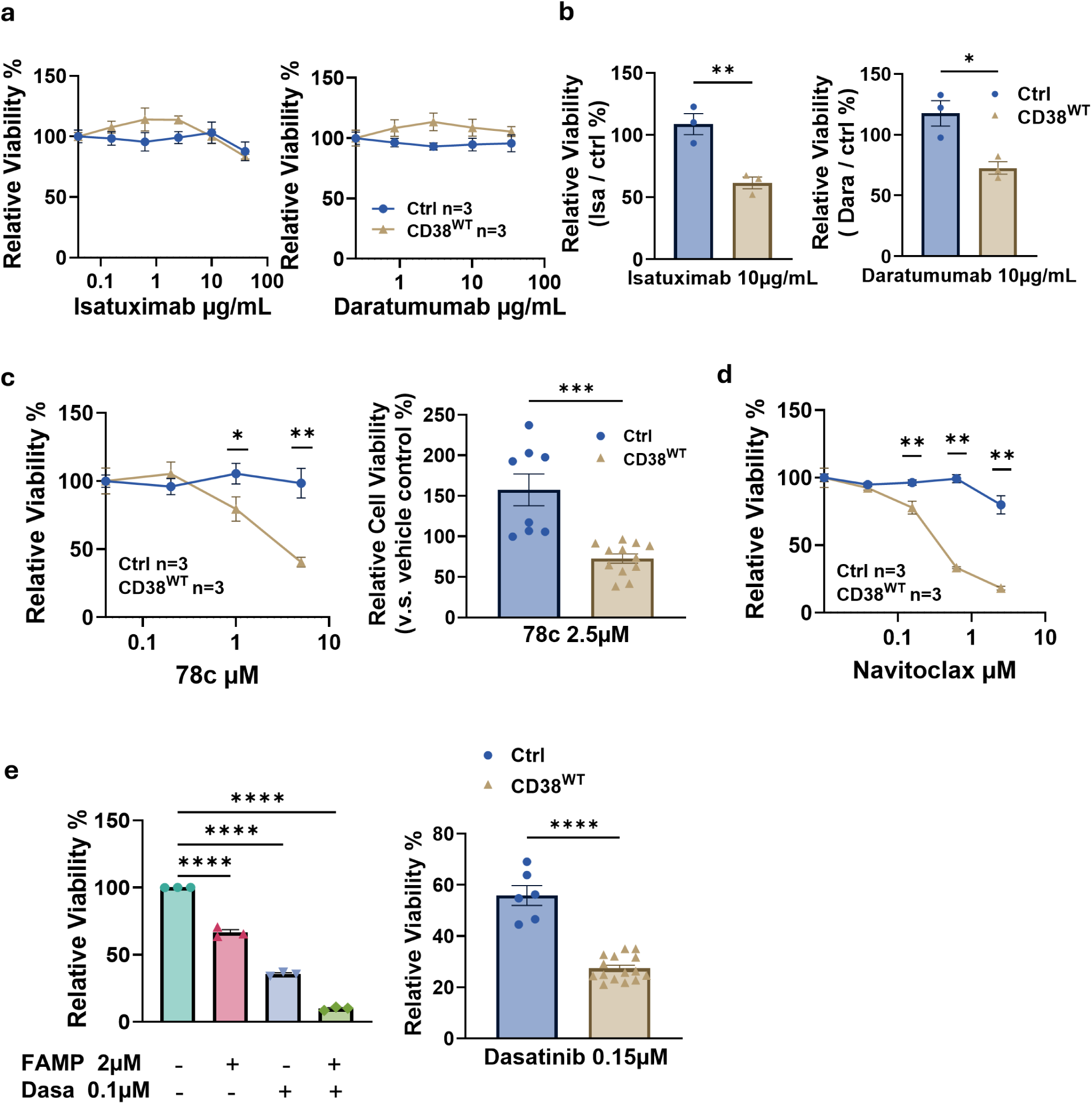
Therapeutic Vulnerabilities of CD38^WT^ MEC-1 Cells to CD38 Inhibition and Senolytic Agents. (a–b) Anti-CD38 monoclonal antibodies daratumumab and isatuximab did not alter MEC-1 cell viability under normoxia after 72 hours of treatment, regardless of CD38 expression (a), but selectively reduced viability of CD38^WT^ cells under hypoxia (2.5% O₂) without affecting vector controls (b). Relative viability compared with vehicle control is shown from representative clones across three independent experiments (n=3; unpaired *t*-test, *: p < 0.05, **: p < 0.01); (c) The small-molecule CD38 inhibitor 78c preferentially reduced viability of CD38^WT^ cells while sparing vector controls after 72 hours of treatment (left panel). Dose–response curves from representative clones are shown (n=3; two-way ANOVA, *: p < 0.05, **: p < 0.01). Right panel showed 78c selectively reduced viability of CD38^WT^ cells under hypoxia (2.5% O₂) without affecting vector controls. Relative viability compared with vehicle control is shown from all tested clones across three independent experiments (unpaired *t*-test, ***: p < 0.001); (d) The senolytic agent navitoclax selectively suppressed CD38^WT^ cell viability compared with vector controls after 72 hours of treatment. Dose– response curves from representative clones are shown (n=3; two-way ANOVA, **: p < 0.01); (e) Dasatinib enhanced fludarabine-associated loss of viability in CD38^WT^ cells (left). Cell viability following 72 hours of treatment with fludarabine (2 µM), dasatinib (0.1 µM), or the combination is shown for a representative CD38^WT^ clone (n=3; unpaired *t*-test, ****: p < 0.0001). When tested alone at 0.15 µM, dasatinib reduced relative viability more strongly in CD38^WT^ cells than in vector controls (right). Data represent the mean of all tested clones from three independent experiments.

### 3.7. Primary CLL Transcriptomic Analyses Support the CD38-Associated Stress-Adaptation Program

To determine whether the molecular phenotype identified in engineered MEC-1 cells is also present in primary CLL, we analyzed three independent single-cell RNA-sequencing cohorts: GSE111014, GSE165087, and GSE259253. We first compared naturally occurring CD38⁺ and CD38⁻ B/CLL cells across the three cohorts. B/CLL cells were identified by lineage-marker expression, and CD38 groups were defined by *CD38* transcript detection, as described in the Materials and Methods. A cohort-balanced summary dot plot showed higher mean normalized *NAMPT* expression and a greater proportion of cells with detectable *NAMPT* transcripts in the CD38⁺ population, with modest differences in *HIF1A* and *CDKN1A* (Figure 7a). Cohort-specific CD38-group dot plots showed the corresponding patterns within each dataset (Supplementary Figure S4a). Dataset-stratified Mantel–Haenszel (MH) analysis showed higher gene-detection odds in CD38⁺ than CD38⁻ cells for *NAMPT* (MH OR 2.26, 95% CI 1.97–2.59, P < 0.001), *HIF1A* (OR 1.40, 95% CI 1.20–1.63, P < 0.001), and *CDKN1A* (OR 1.37, 95% CI 1.13–1.67, P = 0.001; Figure 7b). These findings were consistent with the NAMPT-centered transcriptional pattern identified in the MEC-1 model.

Patient-specific profiles highlighted substantial interpatient heterogeneity across the three cohorts. In GSE111014, the dot plot showed variation in the detection frequency and mean normalized expression of *CD38*, *NAMPT*, *HIF1A*, and *CDKN1A* among the four evaluable patients (Figure 7c), with analogous analyses for GSE165087 and GSE259253 presented in Supplementary Figure S4b. UMAP analysis of GSE111014 further revealed patient-associated transcriptional structure and the spatial distribution of *CD38*-detected and *NAMPT*-expressing cells (Figure 7d). Similar UMAP visualizations for GSE165087 and GSE259253 are provided in Supplementary Figure S4c and S4d, respectively.

To reduce the influence of between-patient differences in genetic background, disease state, treatment history, and transcriptional composition, we next compared CD38⁺ and CD38⁻ cells within the same patient. Patient-stratified analyses showed higher *NAMPT* detection in CD38⁺ than CD38⁻ cells within each cohort: GSE111014 (MH OR 1.95, 95% CI 1.28–2.99, P = 0.002), GSE165087 (MH OR 1.45, 95% CI 1.19–1.77, P < 0.001), and GSE259253 (MH OR 1.82, 95% CI 1.45–2.28, P < 0.001). Across 16 evaluable patients, the overall patient-stratified association remained significant (MH OR 1.63, 95% CI 1.41–1.87, P = 1.25 × 10⁻¹¹), with no statistically detectable heterogeneity among patient-specific odds ratios (Breslow–Day P = 0.893; Figure 7e). Individual-patient odds ratios are shown in Supplementary Figure S4e. Thus, the CD38–NAMPT association persisted after stratification by patient, indicating that it was not solely a consequence of pooling cells from biologically different patients.

As a major ecto-NADase, CD38 catalyzes extracellular NAD⁺ consumption and thereby regulates NAD⁺ availability within the tissue microenvironment [7]. We then asked whether the CD38-associated *NAMPT* expression pattern extended beyond CD38⁺ CLL cells to the CD38⁻ compartment. Accordingly, we assessed whether patients with a larger CD38⁺ CLL-cell compartment also exhibited higher NAMPT detection within their CD38⁻ CLL cells. Across 18 patients, relative CD38⁺ CLL-cell abundance was positively correlated with *NAMPT* detection in the CD38⁻ compartment after within-cohort rank normalization (Spearman ρ = 0.658, P = 0.003, FDR-adjusted P = 0.021; Figure 7f). Thus, patients with a greater relative abundance of CD38⁺ CLL cells also showed higher *NAMPT* detection within their CD38⁻ CLL-cell compartment, consistent with a broader CD38-associated transcriptional pattern across the CLL population.

Collectively, these findings support a CD38-associated NAMPT adaptive program in primary CLL at both the cellular and patient levels. To place these gene-level associations in a broader biological context, pathway-level differences between CD38⁺ and CD38⁻ cells were evaluated. In the 14 patients meeting pathway-analysis criteria, random-effects meta-analysis showed higher pathway scores in CD38⁺ than CD38⁻ cells for Hypoxia (Hedges’ g = 0.15, 95% CI 0.09–0.21, P = 1.80 × 10⁻⁷), PI3K/AKT/mTOR (g = 0.21, 95% CI 0.12–0.29, P = 1.36 × 10⁻⁶), mTORC1 (g = 0.26, 95% CI 0.18–0.33, P = 6.29 × 10⁻¹²), and Glycolysis (g = 0.17, 95% CI 0.12–0.23, P = 6.25 × 10⁻¹¹; Figure 7g). These coordinated transcriptional differences further linked the primary CLL phenotype to the hypoxia- and AKT-associated stress-adaptation program identified experimentally in MEC-1 cells.

Given the limited number of patients available for single-cell analysis, we sought population-level corroboration in larger independent CLL cohorts. In GSE246521 (bulk RNA-seq, n = 350), *NAMPT* expression correlated positively with AKT Score (r = 0.820, P = 2.63 × 10⁻⁸⁶), HYPOXIA Score (r = 0.868, P = 1.08 × 10⁻¹⁰⁷), and *CDKN1A* expression (r = 0.671, P = 3.28 × 10⁻⁴⁷; Figure 7h). Similar associations were observed in the independent GSE58211 microarray cohort (n = 300; Supplementary Figure S4f), in which *NAMPT* expression correlated positively with AKT Score (r = 0.459, P = 4.56 × 10⁻¹⁷), HYPOXIA Score (r = 0.566, P = 8.52 × 10⁻²⁷), and *CDKN1A* expression (r = 0.485, P = 4.53 × 10⁻¹⁹). Together, these independent cohorts provided population-level support for the association of NAMPT expression with AKT- and hypoxia-related transcriptional programs in primary CLL.

**Figure 7.**
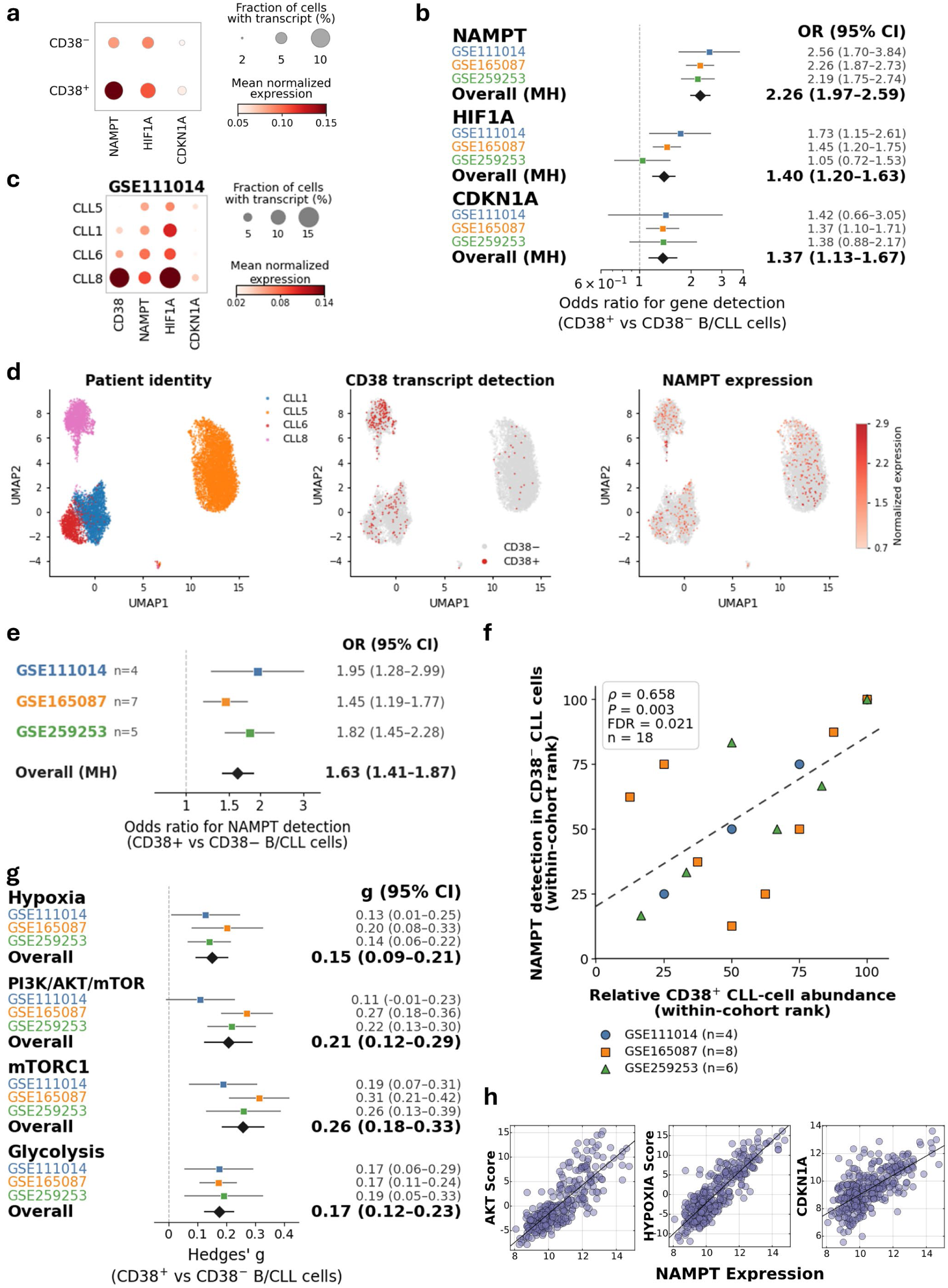
Primary CLL transcriptomic analyses support a CD38-associated NAMPT stress-adaptation program. CD38^+^ and CD38^−^ denote detectable and undetectable CD38 transcripts, respectively, rather than clinical surface-protein categories. (a) Cohort-balanced dot plot of NAMPT, HIF1A, and CDKN1A in CD38+ and CD38− B/CLL cells from GSE111014, GSE165087, and GSE259253. Detection frequency and mean normalized expression were averaged with equal weight across cohorts. Dot size represents detection frequency, and color represents mean normalized expression. (b) Dataset-stratified forest plot of gene-detection odds for NAMPT, HIF1A, and CDKN1A in CD38^+^ versus CD38^−^ B/CLL cells. Colored squares show cohort-specific odds ratios, black diamonds show Mantel–Haenszel pooled estimates, and horizontal lines indicate 95% confidence intervals. Overall estimates were NAMPT, OR 2.26 (95% CI 1.97–2.59, P < 0.001); HIF1A, OR 1.40 (95% CI 1.20–1.63, P < 0.001); and CDKN1A, OR 1.37 (95% CI 1.13–1.67, P = 0.001). (c) Patient-level dot plot for GSE111014 showing CD38, NAMPT, HIF1A, and CDKN1A across the four evaluable patients. Dot size represents the fraction of cells with detectable transcript, and color represents mean normalized expression. (d) UMAP visualization of GSE111014 B/CLL cells showing patient identity, CD38 transcript detection, and NAMPT expression. (e) Within-patient NAMPT-detection association summarized by cohort. Colored squares show cohort-specific Mantel–Haenszel estimates for GSE111014 (n = 4), GSE165087 (n = 7), and GSE259253 (n = 5), the black diamond shows the overall estimate across 16 evaluable patients, and horizontal lines indicate 95% confidence intervals. Cohort-level estimates were GSE111014, OR 1.95 (95% CI 1.28–2.99, P = 0.002); GSE165087, OR 1.45 (95% CI 1.19–1.77, P < 0.001); and GSE259253, OR 1.82 (95% CI 1.45–2.28, P < 0.001). The overall OR was 1.63 (95% CI 1.41–1.87, P = 1.25 × 10^−11^), with no evidence of heterogeneity among patient strata (Breslow–Day P = 0.893). (f) Association between relative CD38^+^ B/CLL-cell abundance and NAMPT detection within the CD38^−^ compartment across 18 patients. Both variables were converted to withincohort ranks before analysis. Each point represents one patient; colors and symbols indicate cohort. Spearman ρ = 0.658, P = 0.003, and FDR-adjusted P = 0.021. The dashed line is a fitted visual guide. (g) Random-effects metaanalysis of pathway-score differences between CD38^+^ and CD38^−^ B/CLL cells. Colored squares show cohort-specific Hedges’ g estimates, black diamonds show pooled cross-cohort estimates, and horizontal lines indicate 95% confidence intervals. Positive Hedges’ g indicates higher pathway scores in CD38^+^ cells. Overall estimates were Hypoxia, g = 0.15 (95% CI 0.09–0.21, P = 1.80 × 10^−7^, I² = 0%); PI3K/AKT/mTOR, g = 0.21 (95% CI 0.12–0.29, P = 1.36 × 10^−6^, I² = 55.6%); mTORC1, g = 0.26 (95% CI 0.18–0.33, P = 6.29 × 10^−12^, I² = 15.6%); and Glycolysis, g = 0.17 (95% CI 0.12–0.23, P = 6.25 × 10^−11^, I² = 0%). (h) Independent bulk RNA-seq validation in GSE246521 (n = 350). NAMPT expression correlated positively with AKT Score (r = 0.820, P = 2.63 × 10^−86^), HYPOXIA Score (r = 0.868, P = 1.08 × 10^−107^), and CDKN1A expression (r = 0.671, P = 3.28 × 10^−47^). Each point represents one patient sample; fitted lines are shown.

## 4. Discussion

Chronic lymphocytic leukemia (CLL) progression and treatment resistance are strongly influenced by metabolic and immunoregulatory signals within the tumor microenvironment [62, 63]. While CD38 has long been recognized as a marker of aggressive disease and inferior survival in CLL, its precise mechanistic role in therapeutic resistance has remained unclear. Our findings indicate that CD38 is not merely a prognostic biomarker, but an active metabolic and signaling regulator that promotes stress adaptation and enhances CLL-cell survival under cytotoxic and hypoxic conditions.

We identify a CD38–NAMPT–AKT–p21 survival axis that links CD38 enzymatic activity to stress adaptation in CLL cells. CD38^WT^ MEC-1 cells, but not catalytically inactive CD38^CI^ cells, showed increased NAMPT expression, enhanced AKT phosphorylation, and accumulation of cytoplasmic p21. NAMPT and AKT signaling have both been implicated in cancer-cell metabolic adaptation [11, 64]. Notably, recent work in Richter transformation of CLL has identified a NAMPT-dependent signaling circuit linked to AKT activation and NAD^+^-regulated deacetylation [65]. Consistent with this mechanistic model, our primary CLL transcriptomic analyses showed that NAMPT expression was associated with CD38 positivity and with AKT- and hypoxia-related transcriptional programs. Together, these findings support a model in which CD38 NADase activity promotes compensatory NAD^+^ salvage and AKT–p21 signaling, thereby enhancing CLL-cell survival under metabolic and cytotoxic stress.

Functionally, CD38-expressing MEC-1 cells displayed markedly reduced apoptosis following fludarabine exposure and under hypoxic stress. This survival advantage depended on CD38 catalytic activity and was attenuated by inhibition of NAMPT or p21. Mechanistically, AKT-mediated phosphorylation provides a plausible explanation for cytoplasmic p21 stabilization, where p21 can interact with procaspase-3 and Bcl-xL to suppress apoptosis [47, 48]. Cytoplasmic p21 has also been linked to enhanced glycolytic adaptation under hypoxic stress [66], consistent with the improved survival of CD38^WT^ cells under low-oxygen conditions. These findings suggest that within hypoxic lymph-node and bone-marrow niches [67], CD38-high CLL cells may be better equipped to persist under metabolic and therapeutic stress. Such stress tolerance could contribute to residual disease and the preferential persistence of CD38-positive subclones during disease progression and treatment. Interestingly, recent work also suggests that CD38 enzymatic activity can influence drug susceptibility in multiple myeloma [68].

Importantly, this stress-adapted phenotype was pharmacologically targetable. NAMPT inhibition with FK866 markedly reduced the viability of CD38^WT^ cells, particularly under stress conditions, while suppression of p21 with UC2288 lowered p21 expression and attenuated resistance to fludarabine and hypoxia. Targeting CD38 enzymatic activity with 78c, as well as treatment with the therapeutic anti-CD38 antibodies daratumumab and isatuximab, also impaired hypoxic adaptation in CD38^WT^ cells. In addition, CD38^WT^ cells showed increased sensitivity to the senolytic agents navitoclax and dasatinib, consistent with acquisition of a stress-tolerant, senescence-like pheno-type. Together, these findings identify several potential therapeutic vulnerabilities in CD38-high CLL, including CD38 blockade, inhibition of NAD^+^ salvage, p21 targeting, and senolytic approaches.

Beyond these tumor-intrinsic effects, CD38 may also influence the broader CLL microenvironment through its ectoenzymatic activity. By regulating extracellular NAD^+^ and NMN availability [7], CD38 can influence metabolic conditions relevant to antitumor immune-cell function [69]. Consistent with this broader role, our primary CLL analysis showed that patients with a larger CD38^+^ CLL-cell compartment also had higher NAMPT detection within their CD38^−^ CLL cells, suggesting that CD38-associated metabolic adaptation may not be confined to the CD38-expressing population. CD38-dependent resistance may be further reinforced by microenvironmental signaling, including CD38 association with CD19/CD81 to augment BCR signaling [22] and CD38/CD31 interactions that support CLL-cell survival [70]. In co-culture systems, CD38 blockade has also been reported to enhance susceptibility to CAR-T-cell-mediated killing [71], supporting further investigation of CD38-directed strategies in combination with cellular immunotherapies.

Although anti-CD38 antibodies have transformed the treatment of multiple myeloma [59], their clinical activity in CLL has been more limited [24, 72]. Our findings suggest that metabolic adaptation through NAMPT-dependent NAD^+^ salvage and downstream AKT–p21 signaling may represent one mechanism by which CD38-high CLL cells maintain survival despite therapeutic stress. Future studies should define more precisely how CD38 NADase activity interfaces with NAMPT, NAD^+^ homeostasis, and AKT–p21 signaling in primary CLL. In vivo studies using CD38-expressing models and primary patient samples will be important for determining whether combined targeting of CD38 and its downstream metabolic dependencies can improve therapeutic responses. Complementary NAD metabolomics and molecular profiling may also help identify biomarkers of CD38-dependent stress adaptation and define the patient populations most likely to benefit from such strategies.

## 5. Conclusions

In conclusion, our findings identify a CD38-dependent stress-adaptation program in CLL in which CD38 NADase activity promotes NAMPT upregulation, AKT activation, and p21-associated survival under hypoxic and cytotoxic stress. Disruption of CD38 catalytic activity or pharmacologic targeting of NAMPT and p21 attenuated this resistant phenotype, revealing potential metabolic vulnerabilities of CD38-high CLL cells. Primary CLL transcriptomic analyses further supported the relevance of the CD38–NAMPT-associated program in patient samples and suggested that CD38-associated metabolic adaptation may also involve the broader CLL microenvironment. Together, these findings provide a mechanistic framework linking CD38-dependent NAD^+^ metabolism to stress tolerance and identify potential therapeutic strategies for targeting treatment-resistant CLL.

## Author Contributions

Conceptualization, B.H., C.C. and E.C.; methodology, B.H. and C.C; validation, B.H., C.C., A.B. and E.C.; formal analysis, B.H., C.C. and A.B.; investigation, B.H. and C.C.; resources, E.B.; data curation, B.H., C.C and E.C.; writing— original draft preparation, B.H. and C.C.; writing—review and editing, G.L., A.P., A.C-K., E.B., Y.H. and E.C.; supervision, C.C. and E.C.; funding acquisition, A.C-K. and E.C. All authors have read and agreed to the published version of the manuscript.

## Institutional Review Board Statement

Not applicable.

## Informed Consent Statement

Not applicable.

## Data Availability Statement

Publicly available datasets analyzed in this study are available through the NCBI Gene Expression Omnibus under accession numbers GSE58211, GSE246521, GSE111014, GSE165087, and GSE259253. Additional processed analysis tables generated for the revised study are available from the corresponding author upon reasonable request.

## Abbreviations

The following abbreviations are used in this manuscript:

CLL: Chronic lymphocytic leukemia
NAD: Nicotinamide adenine dinucleotide
NAMPT: Nicotinamide phosphoribosyltransferase
CD38: Cluster of Differentiation 38
Cadpr: Cyclic ADP-ribose
PARP: Poly (ADP-ribose) polymerase
BCR: B cell receptor
HIF1A: Hypoxia-inducible factor 1-alpha
VEGFA: Vascular Endothelial Growth Factor A
LDHA: Lactate Dehydrogenase A
GLUT1: Glucose transporter type 1
Bcl-xL: B-cell lymphoma-extra large
BCL-2: B-cell lymphoma 2
WT: Wild type
CI: CD38 Catalytically inactive

