## Supplementary figures and images for "CD38–NAMPT–p21 signaling promotes stress adaptation in CLL"

### SUPPLENENTAL FIGURES

#### Slide 1
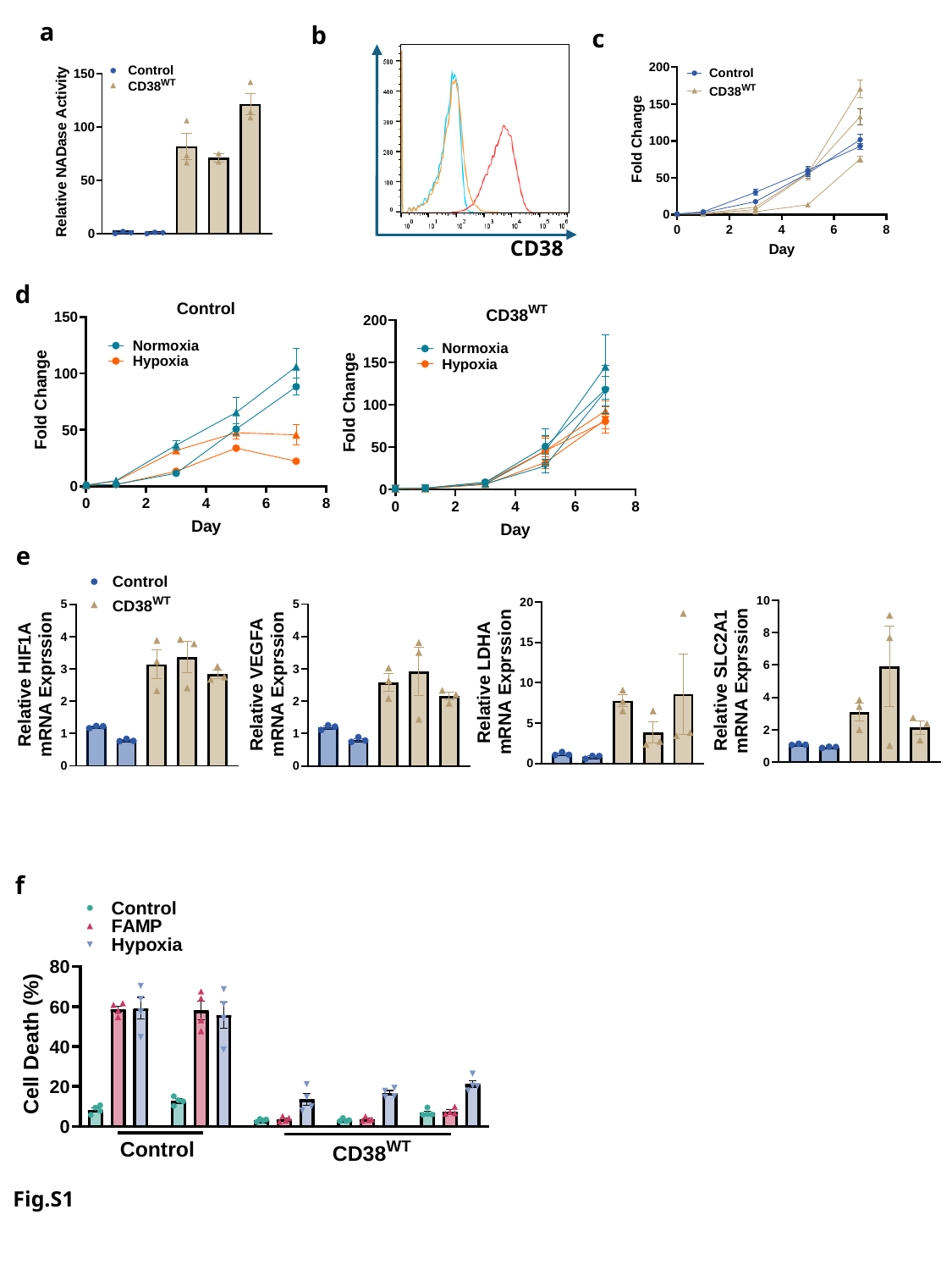

a
b
c
CD38
d
e
f
Fig.S1

#### Slide 2
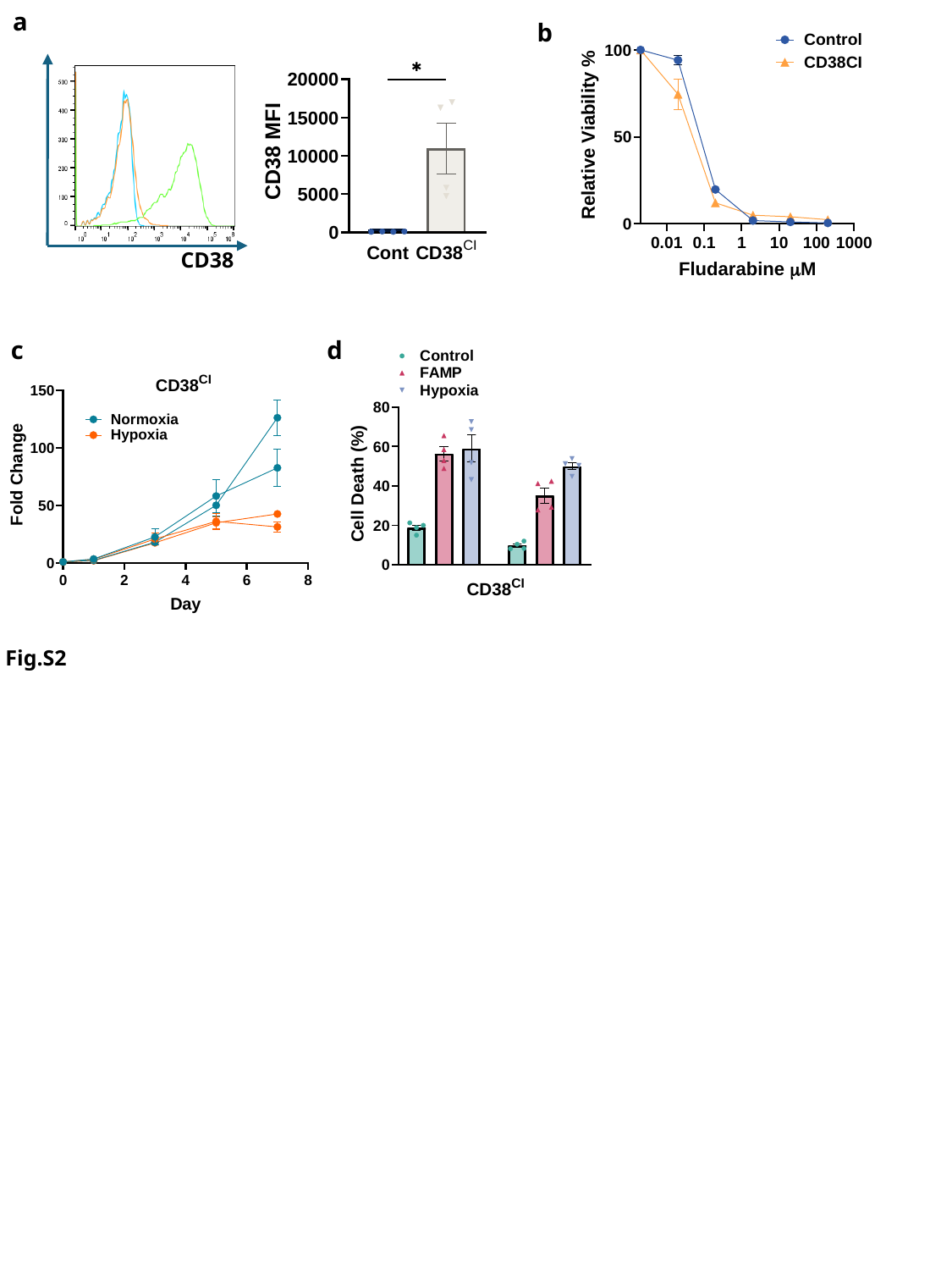

a
b
CD38
c
d
Fig.S2

#### Slide 3
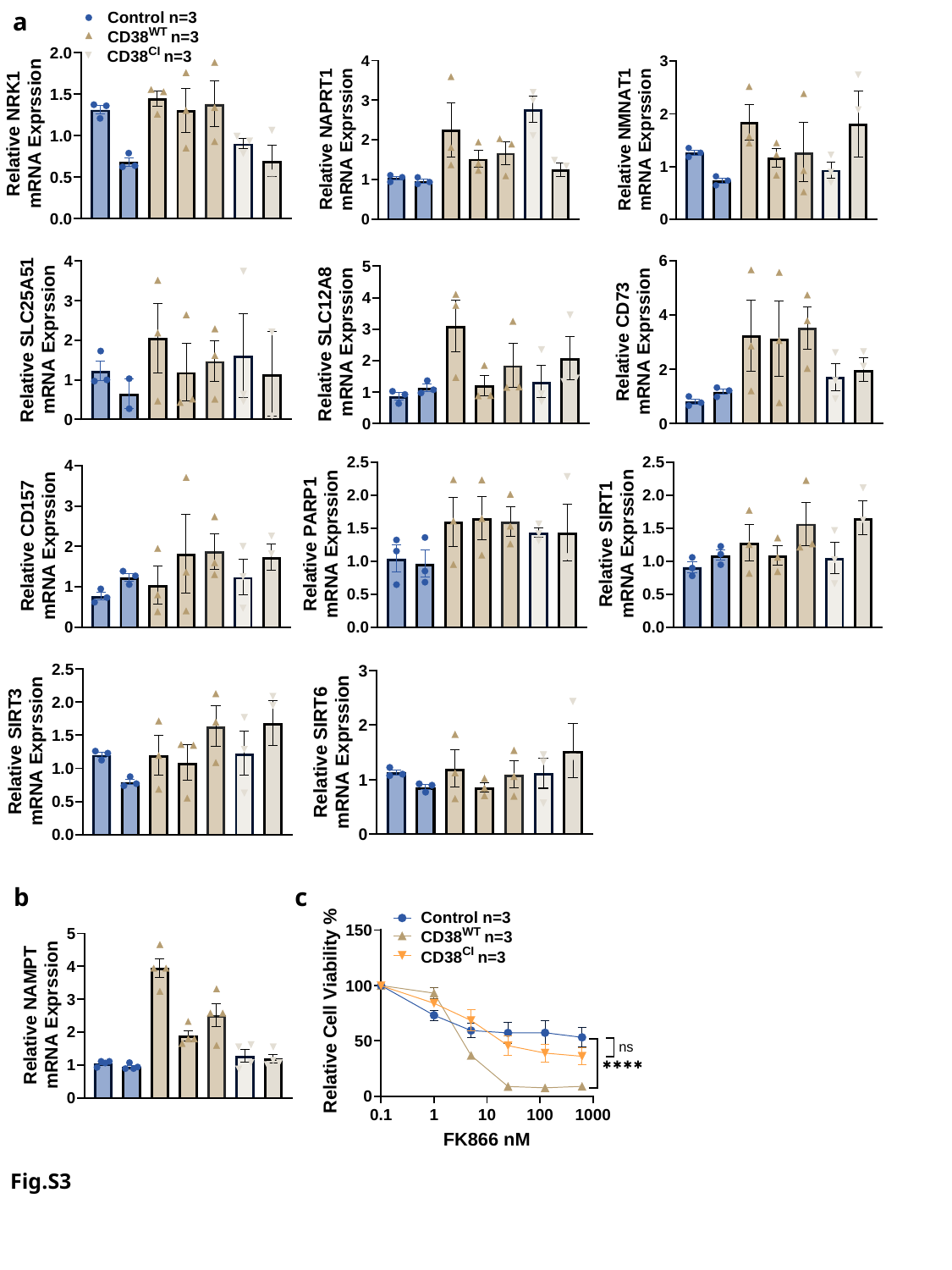

a
c
b
Fig.S3

#### Slide 4
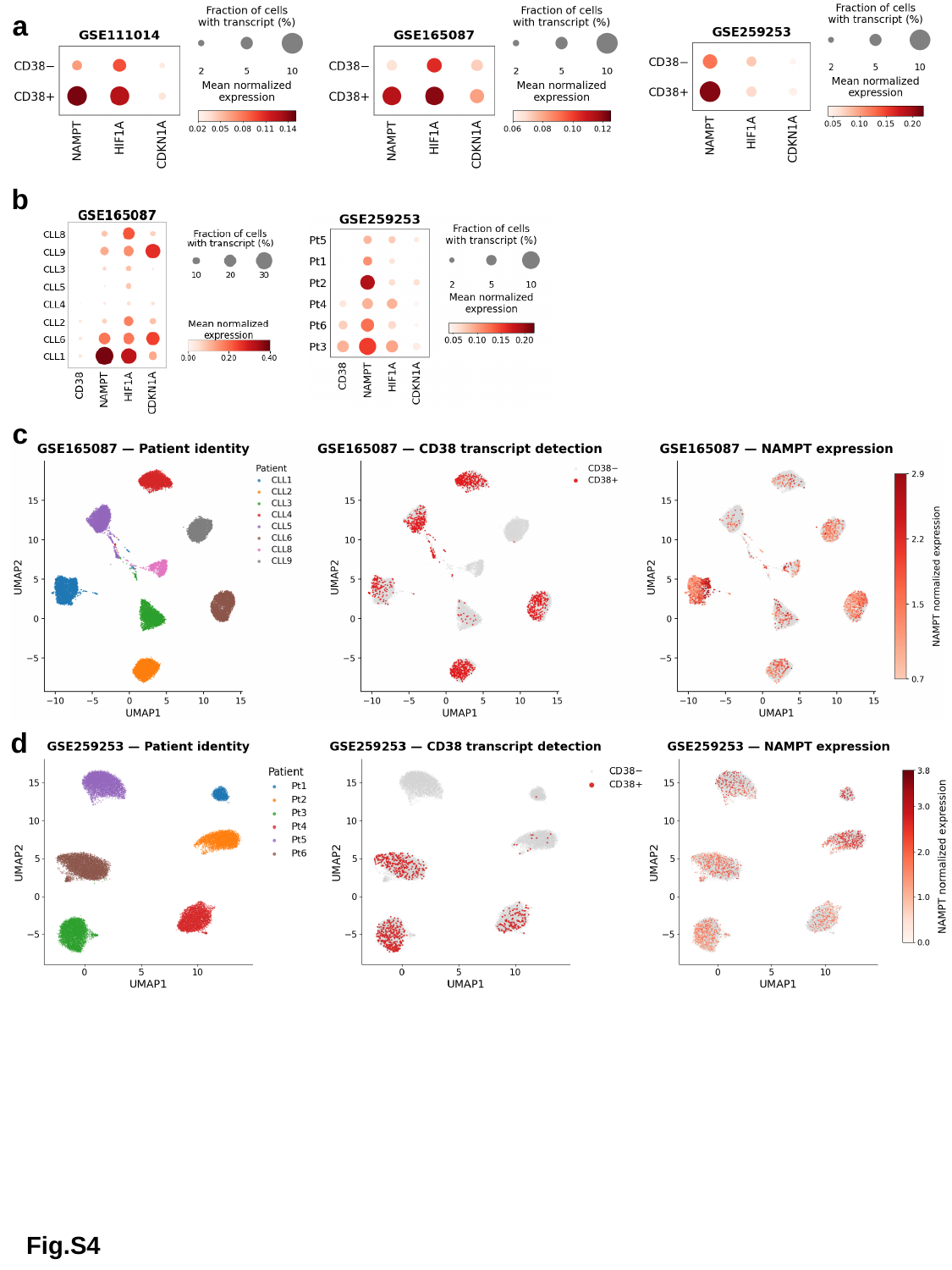

a
b
c
d
Fig.S4

#### Slide 5
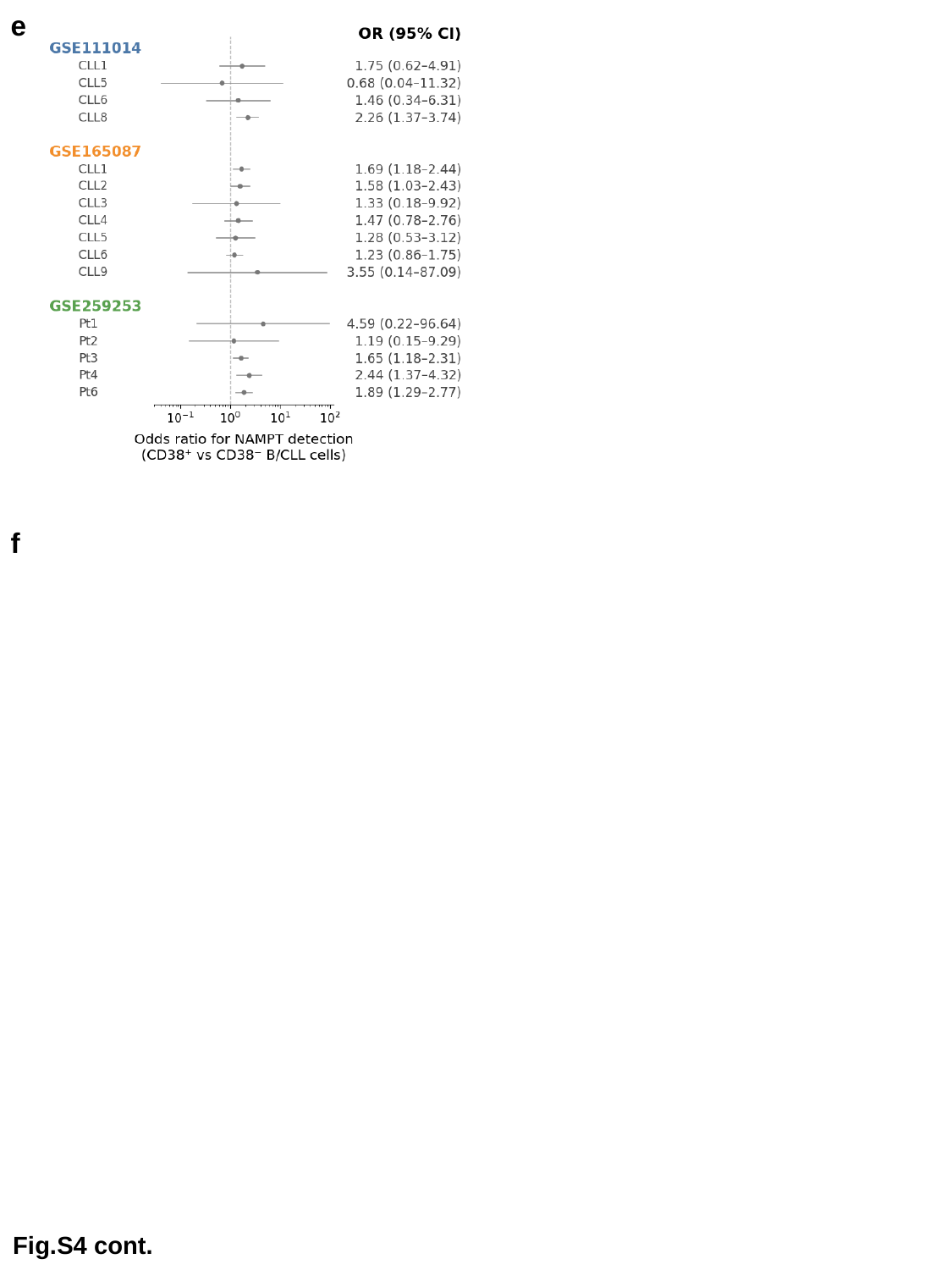

e
f
Fig.S4 cont.
